# Skin-conformal electronics for wearable electrogastrography monitoring

**DOI:** 10.1101/2025.10.14.682324

**Authors:** Liuxi Xing, Yulu Cai, Yunnuo Zhang, Vittorio Mottini, Linux Heller, Jinxing Li

**Affiliations:** Department of Biomedical Engineering and Institute for Quantitative Health Science & Engineering, Michigan State University, East Lansing, MI, USA; Department of Chemical Engineering and Material Science, Michigan State University, East Lansing, MI, USA; Wallace H. Coulter Department of Biomedical Engineering, Emory University and Georgia Institute of Technology, Atlanta, GA, USA; Department of Biomedical Engineering and Institute for Quantitative Health Science & Engineering, Department of Chemical Engineering and Material Science, Department of Electrical and Computer Engineering and Neuroscience Program, Michigan State University, East Lansing, MI, USA

**Keywords:** Soft electronics, Wearable sensors, Electrogastrogram (EGG), Electrocardiogram (ECG)

## Abstract

Electrogastrography (EGG), a non-invasive method for measuring gastric myoelectrical activity, offers valuable insights into gastrointestinal motility and functional disorders such as gastroparesis and functional dyspepsia. Despite its diagnostic potential, the clinical adoption of EGG has been limited due to its reliance on rigid electrodes and bulky instrumentation, which leads to motion artifacts and poor signal quality, and ultimately reduces patient comfort and restricting data collection to short-duration, stationary settings. To address these limitations, we present FlexEGG, a skin-conformal, flexible electronic system engineered for high-fidelity EGG monitoring in both clinical and real-world environments. The device incorporates a soft, stretchable electrode array specifically designed for the abdominal surface, utilizing a hybrid stretchable conductor composed of conductive polymer, silver nanowires (AgNWs), and polyurethane elastomer, which leads to good skin contact, signal stability, and mechanical conformability. A custom low-noise analog front-end and digital signal processing pipeline enables reliable acquisition of low-frequency, low-amplitude gastric slow waves. Additionally, FlexEGG supports simultaneous electrocardiogram (ECG) measurement, potentially facilitating integrated gut–heart axis monitoring for broader physiological assessment. In this study, we describe the design, implementation, and validation of FlexEGG in multi-channel, long-duration EGG and ECG recordings. Our findings demonstrate its potential as a wearable, non-invasive tool for continuous gastrointestinal electrophysiology monitoring, enabling new opportunities for diagnosing and managing digestive disorders in everyday settings.

## I. INTRODUCTION

Gastrointestinal motility disorders (GMDs), such as gastroparesis, functional dyspepsia, and irritable bowel syndrome (IBS), affect a significant portion of the population and are often associated with chronic pain, bloating, nausea, and impaired digestion[1], [2]. These conditions are especially common in aging individuals and patients with diabetes, Parkinson’s disease, or other autonomic dysfunctions[2]–[5]. However, diagnosis remains challenging due to the intermittent and nonspecific nature of symptoms[1], [6], [7]. There is a critical need for non-invasive tools that enable objective, continuous monitoring of gastric physiology to support early diagnosis, treatment optimization, and long-term disease management.

Electrogastrography (EGG) is a non-invasive technique for recording gastric myoelectrical activity via surface electrodes placed on the abdomen[8]–[11]. EGG offers a promising window into gastric slow-wave activity, which plays a central role in coordinating gastric motility[12]–[14]. Abnormal slow-wave patterns, such as bradygastria, tachygastria, and dysrhythmias, have been correlated with various motility disorders[15], [16]. Additionally, EGG may provide biomarkers for understanding gut–brain signaling and autonomic regulation[17]–[20]. However, its clinical utility has been limited by technical challenges: the signals are of low amplitude and frequency (typically 0.05–0.2 Hz), and are easily contaminated by noise from respiration, cardiac activity, and motion artifacts[14]. Conventional EGG systems rely on rigid electrodes and cumbersome setups, limiting their suitability for ambulatory use and real-world data collection[21]–[26].

Translating EGG into wearable health monitoring devices presents several obstacles. First, multi-channel signal acquisition typically requires multiple wired electrodes, which can be uncomfortable and restrict user mobility. Second, reliable long-term recording demands stable skin-electrode contact and robust signal quality, which are difficult to maintain using conventional dry or gel-based electrodes especially during daily activities. Third, the real-time processing of low-frequency, low-amplitude signals requires careful noise filtering and motion artifact suppression, which often conflict with the design goals of miniaturization and low power consumption.

Recent advances in skin-conformal electronics have enabled the development of wearable systems that combine high mechanical compliance, biocompatibility, and signal fidelity[27]–[36]. These systems have shown promise in applications such as EEG, EMG, and ECG[37]–[50]. However, reliable EGG monitoring in dynamic, real-life settings has been scarcely explored. Gastric slow-wave signals have intrinsically low frequencies and small amplitudes, and their acquisition is further complicated by inter-individual variations in skin hydration, texture, and elasticity[51]. Commercial wet electrodes, while widely used, often rely on conductive gels that can dry during extended recordings, leading to increased impedance and degraded signal quality. Such variability often leads to elevated impedance, inconsistent electrode contact, and unstable signal quality, posing unique challenges for non-invasive gastric electrophysiology [52]–[54].

Building upon the advanced soft bioelectronics, we sought to extend these benefits to non-invasive gastric electrophysiology. Here, we present FlexEGG, as shown in Fig. 1(a), a skin-conformal, flexible bioelectronic platform specifically designed for high-fidelity, multi-channel EGG monitoring, with the additional capability of simultaneous electrocardiography (ECG) acquisition. As shown in Fig. 1(b-c), the system features a soft, stretchable electrode array composed of poly(3,4-ethylenedioxythiophene)-poly(styrenesulfonate) (PEDOT:PSS), silver nanowires (AgNWs), waterborne polyurethanes (WPU), ionic liquid, and dimethyl sulfoxide (DMSO) additives to achieve high stretchability, conductivity, and robust skin adhesion. Glycerol, a biocompatible humectant, can enhance PEDOT:PSS chain mobility, prevent cracking, and increase skin-electrode adhesion through hydrogen bonding[55]–[59]. The ionic liquid and silver nanowires further tune the impedance by promoting a PEDOT rich region and polymer chain alignment[60]–[62]. WPU provides more elasticity ensuring conformal contact with various skin types[63]. The incorporation of all components significantly increases the overall performance of the electrodes used for high fidelity recording. The compact, wearable hardware integrates a low-power analog front-end (TI AD8232) and a Nordic nRF52840 system-on-chip for wireless data transmission, enabling real-time monitoring in everyday environments, as shown in Fig. 1(d).

**Fig. 1.**
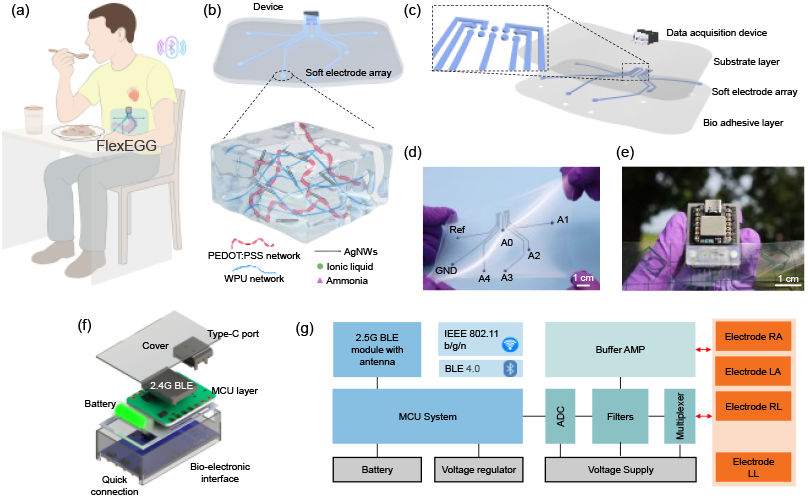
Soft multichannel FlexEGG monitoring system. (a) Schematic of the wearable EGG system applied to the human body, consisting of a soft electrode array and a detachable acquisition device. (b) A detailed illustration of the multichannel soft electrode array and material compositions. (c) 3D-rendered structure of the multilayer device. (d)(e) Optical images of the FlexEGG monitoring system, including (d) multichannel electrode array and (e) device hardware. (f) The framework design of the multiple-channel signal acquisition device shows the main components, including the bio interface, the MCU for data processing, and the wireless module for data transmission. (g) The electrical system of the wearable device.

The main contributions of this work are: The design and implementation of FlexEGG, a flexible, multi-channel wearable platform for continuous EGG and ECG monitoring; material and system-level optimization to achieve stable skin contact, high signal quality, and low power consumption suitable for long-term use; validation of FlexEGG in multi-site recordings during routine activities, demonstrating its potential for a non-invasive, real-world assessment of gastrointestinal electrophysiology. This paper details the device architecture, soft electrode material development, signal processing strategies, and performance evaluations, establishing FlexEGG as a promising tool for advancing unobtrusive gut monitoring and enabling new insights into gut–heart interactions.

## II. MATERIALS AND METHODS

### A. System Setup

The FlexEGG comprises a skin-conformal multiple-channel soft electrode array and custom-made hardware to monitor the epidermal electrophysiological signals, and a low-power multiple-channel bioelectric signal acquisition device with wireless data transmitting function, as illustrated in Fig. 1(a). Fig. 1(b) presents a detailed illustration of the multichannel soft electrode array and material compositions. The electrode array is fabricated using a solution-processable composite material consisting of PEDOT:PSS and waterborne polyurethane (WPU), which is mixed to combine their respective electrical and mechanical properties. To further enhance the conductivity, elasticity, adhesion, and stability, silver nanowires, DMSO and ionic liquids (EMIM:ESO_4_) are introduced as additives. Specifically, PEDOT:PSS serves as the primary conductive polymer, providing both electronic and ionic conduction pathways[64]. Silver nanowires, as highly conductive electronic pathways, bridge PEDOT-rich domains and facility long-range charge transport. Glycerol, a biocompatible humectant, improves chain mobility of PEDOT, suppresses microcrack formation under strain, and increases skin adhesion through hydrogen bonding[65]. Ionic liquid additives further tune impedance by disrupting the hydrogen-bonding network within PSS, promoting PEDOT chain alignment, achieving simultaneously reducing the modulus and enhancing stretchability[66]. Waterborne polyurethane acts as an elastic matrix that maintains mechanical integrity and conformal skin contact under deformation[63]. In addition, DMSO acts as a hygroscopic additive that retains moisture within the composite and promotes PEDOT:PSS chain rearrangement, thereby enhancing conductivity. This organic–inorganic mixture is to balance the multidimensional needs of flexibility, adhesion, conformability, and stability suited to low-frequency EGG in daily use. The resulting ultrathin (∼100 µm) composite electrodes are non-toxic and show no observable skin irritation during continuous wear, as confirmed in biocompatibility tests (Supplementary Fig. S1). The resulting conductive nanocomposite exhibits excellent conformability and low interfacial impedance, ensuring high signal quality during motion and over extended periods. This stretchable electrode array was then interfaced with the exposed contacts on the printed circuit board (PCB) of the device using POGO pin connector. The device is designed as a portable, multi-channel bioelectrical signal acquisition unit that interfaces with the stretchable array to acquire skin potential across multiple channels and reconstructs multi-lead EGG and ECG signals through embedded processing algorithms. Fig. 1(c) presents a 3D rendering of the entire wearable device, highlighting its core components: the bio-adhesive layer, the soft electrode array, the substrate, and the data acquisition device.

Fig. 1(d) and (e) shows the photos of the multiple-channel FlexEGG stretchable electrode array and the device. The resolution and pattern fidelity demonstrate compatibility with laser ablation and liquid processing techniques. The system features 7 electrodes (A0 to A4, reference electrode, and ground electrode), allowing for a configuration that approximates a 4-channel EEG and a 5-channel ECG test. Specifically, it includes direct measurement of the ECG signal, while the EGG signal can be derived mathematically from the recorded electrophysiological data. Using a 3-electrode measurement setup, 5-channel ECG signals are acquired from electrode groupings consisting of a ground electrode, a reference electrode, and a signal electrode (Ax), where Ax refers to electrodes A0 through A4. To calculate the four-channel EGG signals (Ch1 to Ch4), a differential operation is performed such that Chx = Ax – A0, where Ax corresponds to electrodes A1 through A4. The framework of the device is shown in Fig. 1(f), which illustrates the main components, including data acquisition, signal processing, and wireless transmission. The device integrates a wireless transmission module, a microcontroller unit (MCU), and an analog front-end (AFE) into a compact, battery-powered platform optimized for mobility and low power consumption. The electrical system configuration is shown in Fig. 1(g). In our implementation, BLE scanning and data parsing were achieved using the Bluefruit protocol, with communication to a PC via the Bleak Python API for universal beacon scanning. This protocol can also be readily deployed on smartphones with Bluetooth and a modern operating system.

### B. Fabrication Process and Structural Validation

To achieve maximum adaptability across different skin types, we developed a solution-processable conductive polymer composite for fabricating the soft electrode array. This array is designed to be stretchable and soft, and have low skin-electrode impedance, enabling a stable and conformal interface with dynamic, textured, and deforming skin surfaces. The core conductive material consists of PEDOT:PSS, WPU and silver nanowires, which together form a percolated conductive network that combines high electrical conductivity with mechanical stretchability, as shown in Fig. 2(a). This blend strategy leverages the processability of PEDOT:PSS while utilizing WPU’s elasticity, resulting in a soft and conformal material system that is suitable for large-area fabrication via liquid-phase processing. To enhance the mechanical and electrical properties, 1-ethyl-3-methylimidazolium ethyl sulfate (EMIM:ESO_4_) as an ionic liquid and AgNWs were incorporated. A small amount of ammonia was added to prevent aggregation of the anionic WPU, which could otherwise be induced by the acidic sulfonate groups in PEDOT:PSS[67]. Dual-step post-processing further enhances stability and stretchability. In the first step, slow water evaporation leads to a partial assembly of a polymer and elastic WPU matrix. The rapid DMSO evaporation further facilitates polymer chain rearrangement and connection between silver nanowires.

**Fig. 2.**
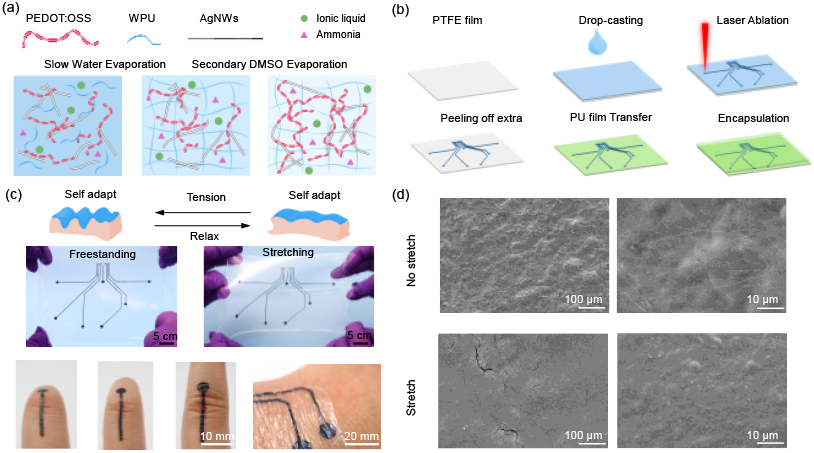
Fabrication process and materials characterization of the stretchable electrode array. (a) Schematic illustration of the conductive composite network and formulation. (b) Stepwise fabrication process of the soft electrode array via drop-casting, laser ablation, transfer, and encapsulation. (c) Photographs of the composite electrode in freestanding and stretching conditions, as well as conformal attachment to skin. (d) SEM images of the electrode surface morphology before and after 20% strain.

The fabrication process shown in Fig. 2(b) begins with drop casting the conductive mixture onto a PTFE film. After dual-step drying, the film is patterned by CO_2_ laser ablation based on the designed multiple-channel pattern. The patterned film is then transferred to a soft PU substrate and encapsulated with another PU film, which also exposes the electrode area. Fig 2(c) confirms the mechanical adaptability of the electrode under various conditions, including freestanding, bending, stretching, and skin conforming. SEM imaging (Fig. 2(d)) reveals the surface morphology of the composite film before and after stretching. Compared to the unstretched state, only minor cracks were observed.

### C. Electrical Stability Test and Formula Optimization

To evaluate the mechanical and electrical performance of the conductive polymer composite, we characterized its electromechanical stretchability and cyclic durability. A 10 mm × 5 mm rectangular sample of the stretchable conductor was removed from the PTFE film for material testing. Electromechanical stretchability was assessed by measuring the relative change in resistance (R/R_0_) under tensile strain, for both the free-standing composite and the composite transferred onto an elastic polyurethane (PU) substrate. Fig. 3(a) shows the variation in resistance as a function of strain. While our materials characterization demonstrates survivability up to 350% tensile strain, ambulatory EGG on the abdomen typically imposes ≤15% strain during daily activities (relaxed/deep breathing, walking, posture changes). Accordingly, our durability benchmarks target ≤20% strain to cover this envelope, and the recorded stability under 20% cyclic loading (Fig. 3b,d) supports reliable operation within physiologic motion ranges. To further assess strain sensitivity under conditions resembling daily use, we measured the resistance– strain behavior of the composite transferred onto a PU substrate, mimicking direct skin attachment. As shown in Fig. 3(c), the conductor again demonstrated a stable response under 20% uniaxial strain. Fig. 3(d) presents the corresponding cyclic durability on the PU substrate, where the resistance change remained approximately around 1, indicating excellent mechanical and electrical stability.

**Fig. 3.**
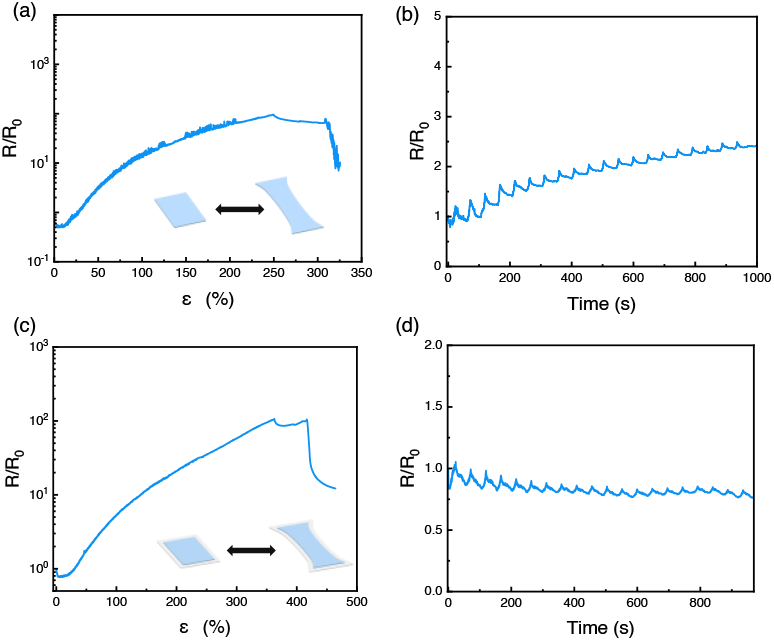
Electrical stretchability of the electrode. (a) Resistance– strain behavior of the free-standing stretchable conductor. (b) Cyclic resistance changes under repeated stretching (20% strain). (c) Resistance–strain behavior of the stretchable electrode transferred onto an elastic PU substrate. (d) Cyclic resistance response of the transferred electrode under 20% cyclic strain.

### D. Thermal Stability of Composite Electrodes

To study the thermal stability of the optimized composite electrode, we examined its temperature-dependent electrical impedance. As shown in Fig. 4(a), the resistance was measured across a temperature range from 23 °C to 60 °C, revealing a relatively low impedance variation. This small change is beneficial for maintaining signal fidelity during real-world use. The frequency-dependent impedance behavior at relevant frequencies (10 Hz, 100 Hz, and 1000 Hz) was also measured and showed consistent results, as illustrated in Fig. 4(b). The tested temperature range (23–60 °C) was chosen to provide design margin, while within the physiological range of 30–40 °C the impedance variation is <20% and does not affect recording fidelity of low-frequency gastric signals.

**Fig. 4.**
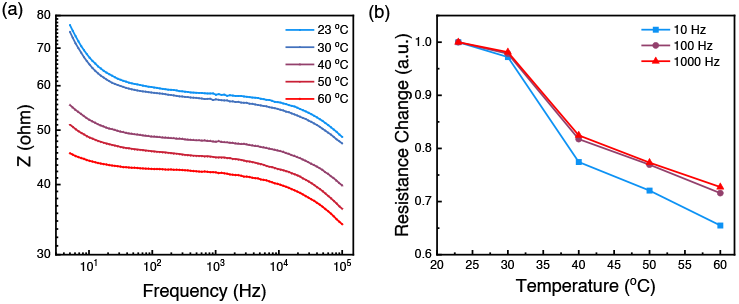
Thermal stability of electrode. (a-d) Frequency-dependent impedance at various temperatures (23°C, 30°C, 40°C, 50°C, 60°C). (b) Temperature-dependent impedance with selected frequency conditions (10 Hz, 100 Hz, 1000 Hz).

## III. RESULTS AND DISCUSS

### A. Multiple Channel ECG Data Test

Gastric slow waves are typically characterized by low amplitude (<500 μV), low frequency (<0.05 Hz or <3 cycles per minute), and slow propagation from the corpus to the pylorus. Compared to ECG, EGG signals place higher demands on the front-end circuit, requiring high input impedance, high common-mode rejection ratio (CMRR), low-frequency response (down to <0.05 Hz), and low-noise amplification.

Here, we design a novel differential measurement system that incorporates two AD8232 chips. The differential output is used to represent the EGG signal, while the independent channels can also capture ECG information. The core of the proposed system is the modified AD8232 analog front-end, tailored for short-term abdominal surface recording in seated or semi-recumbent positions. It includes an instrumentation amplifier, configurable high-pass and low-pass filters, and a right-leg drive (RLD) amplifier for common-mode feedback. Unlike its conventional use for ECG, the filter circuits were reconfigured to accommodate low-frequency gastric signals by increasing the coupling capacitor and feedback resistor. Additionally, the RLD electrode was repositioned to the lower right abdomen far from the stomach, where the minimal (∼0 mV) ECG signal intensity helps to reduce interference.

The microcontroller samples the signal at 1 kHz, providing ample margin for oversampling and later down sampling. Multi-channel skin electrophysiological signals are captured by the system. The system operates in three modes: standby (idle state), monitor (continuous data acquisition), and high-power mode (12-lead ECG acquisition, processing, and Bluetooth data transmission). The device is powered by a Li-polymer battery (nominal voltage: 3.7 V, fully charged at 4.2 V), regulated through a linear voltage regulator. An onboard USB-C port is included for data transfer, firmware updates, and battery charging. Battery capacity can be adjusted depending on application and size constraints. Ethical approval for wearable electrophysiological monitoring was obtained from the IRB Committee at Michigan State University (Approval No. STUDY00011249). For daily wearable monitoring (monitor and alarm mode), the device consumes an average current of ∼200 μA, allowing for over one week of continuous operation with a standard commercial LiPo battery (1C–3.7 V–190 mAh; Grepow Inc.).

The contact area of the electrode array is circular, with a diameter of 5 mm, following the standard design for wearable dry electrodes used in epidermal biopotential detection. The electrodes yield high-quality ECG signals with well-defined PQRST morphologies and a peak-to-peak QRS amplitude of approximately 1.53 mV, comparable to those obtained using standard ECG systems. The differential signal between electrode A0 (see Fig. 1) and the other electrodes represents the raw EGG signal. Fig. 5(a) illustrates the data processing pipeline using channel A0 and A1 as an example.

**Fig. 5.**
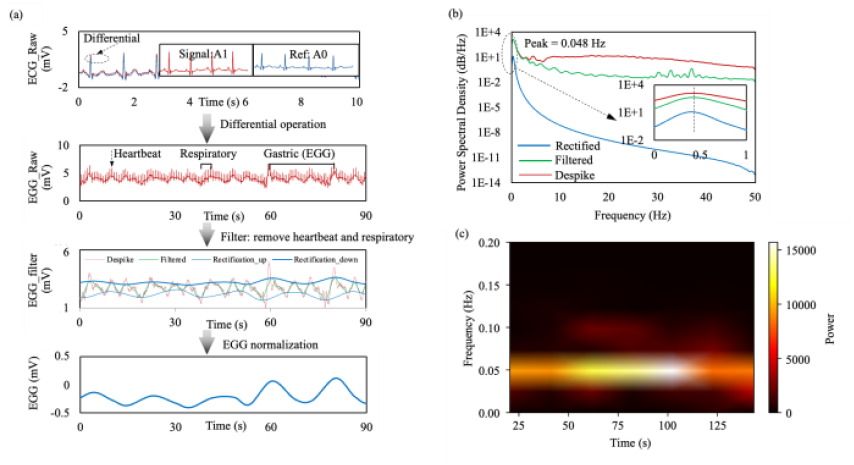
The data processing of single channel ECG/EGG signals. (a) Entire signal processing for extracting gastric slow wave components from raw signals, including the differential operation of two ECG signals A1 and A0, different components recognition (low-frequency gastric slow waves, respiratory artifacts, and heartbeat), and multi-step filtering approach to get the final EGG wave. (b)(c) The frequency-domain analysis of processed EGG signals using (b) PSD analysis and (c) time-frequency spectrogram. It shows the frequency characteristic of EGG wave and other accompanying signal.

The raw EGG signal contains multiple overlapping physiological components, including low-frequency gastric slow waves (∼0.05 Hz or ∼3 cycles per minute), respiratory artifacts (typically 0.2–0.4 Hz), and cardiac interference (>1 Hz). To isolate the gastric component, a two-step filtering approach is employed. First, a spike detection algorithm is used to remove heartbeat-induced spikes. Then, a bandpass filter is applied to isolate the gastric frequency range (typically 0.015– 0.15 Hz in this study), effectively suppressing respiratory and cardiac artifacts while preserving the underlying gastric waveform.

A Savitzky-Golay filter is utilized for smoothing due to its ability to perform local polynomial regression while preserving signal amplitude, shape, and peak timing [68]. This is particularly important for periodic biological signals like gastric slow waves, where conventional IIR filters such as Butterworth or Chebyshev may introduce phase distortion or attenuate important features in the ultra-low frequency band (<0.1 Hz), compromising temporal fidelity and rhythm analysis accuracy[68], [69]. The processed signal reveals a clear, periodic oscillatory pattern consistent with physiological gastric slow waves, enabling quantitative analysis of amplitude, frequency, and propagation dynamics. For verify the performance of our proposed device, a comparison experiment between the FlexECG system and a high-performance wearable commercial holter is launched, supporting the effectiveness of our system in acquiring reliable biopotential signals (Supplementary Fig. S2).

The power spectral density (PSD) plot of the EGG signal in Fig. 5(b) reveals its dominant frequency components after signal preprocessing. A prominent spectral peak is observed at approximately 0.048 Hz, corresponding to a gastric slow-wave rhythm of ∼2.88 cycles per minute (cpm), which aligns with typical human gastric pacemaker activity. The inset highlights the ultra-low-frequency range (0–1 Hz), where the majority of EGG signal energy is concentrated. Higher-frequency components (>0.2 Hz), primarily associated with respiration and cardiac activity, are significantly attenuated following the filtering and smoothing steps. A log–log scale is used in the PSD plot to enhance separation between physiological signals and broadband noise.

Fig. 5(c) presents the time–frequency spectrogram of the filtered EGG signal. A consistent energy band is observed throughout the entire recording period, exhibiting minimal frequency drift. This temporal coherence is indicative of normal, well-entrained gastric motility and highlights the system’s capability for reliably detecting physiological gastric rhythms.

Fig. 6(a) shows the electrode placement schematic for a 4-channel EGG recording. Four recording electrodes (A1–A4) were positioned along the stomach axis, following a 45° oblique line extending from the left hypochondrium to the right lower abdomen. The contrast electrode (A0) was placed below the xiphoid process. The reference and ground electrodes were positioned laterally on the upper and lower right quadrants, respectively. This configuration enables the detection of ECG signals from A0 to A4 and supports the calculation of the spatiotemporal propagation of gastric slow waves using a differential algorithm.

**Fig. 6.**
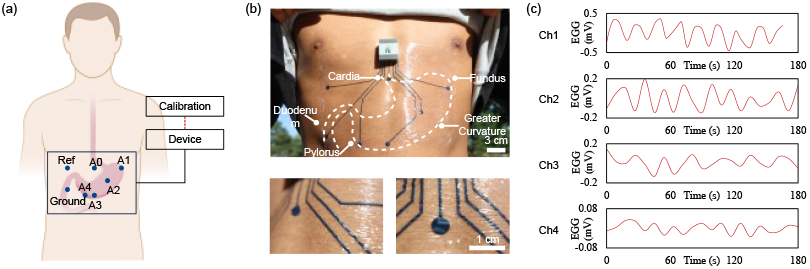
The four-channel EGG setup and representative signals. (a) The position of 7 electrodes, including reference, ground, and A0 to A4. (b) Picture of ECG/EGG soft electrode array and device wore by a participant. (c) Typical four channel EGG signal over 3 minutes.

Using A0 as a common reference, the signals captured by A1– A4 reflect EGG activity from the proximal to distal regions of the stomach. The actual device and soft electrode array worn by a participant are shown in Fig. 6(b). Fig. 6(c) displays the calculated EGG signals from the four channels (Ch1–Ch4) over a 3-minute interval. Each channel exhibits a rhythmic, low-frequency oscillation consistent with gastric slow wave activity (∼3 cycles per minute). Notably, a gradual phase delay is observed from Ch1 to Ch4, indicating the directional propagation of gastric electrical activity from the proximal to the distal stomach.

### B. Continuous Monitoring

The EGG system was worn by a participant for long-term monitoring, as shown in Fig. 7(a). During the 12-hour test period (from 9 a.m. to 9 p.m.), the participant engaged in typical daily activities, including sitting, resting, and indoor work. Although some noise accumulation was observed over time, the EGG system maintained several key advantages: a thin and lightweight design, high signal stability during movement, and comfort while wearing that did not interfere with normal behavior. The entire system operated wirelessly, enabling non-invasive, continuous, and real-time monitoring in real-life settings. Additionally, the device includes ports for auxiliary analog-to-digital converter (ADC) input, allowing future integration of thermistors and photoplethysmography (PPG) sensors for onboard temperature and heart rate measurement.

**Fig. 7.**
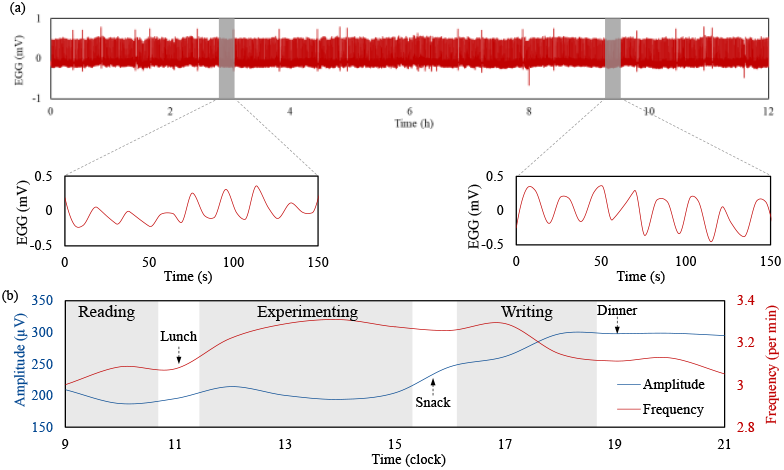
The continous monitoring in multiple working mode. (a) A 12 h test on channel A1. During the testing, the participant is requested to wear the device and engage in daily activities while the EGG signal from channel A1 is wirelessly collected without interrupting the device and measurements. (b) Frequency and amplitude analysis of gastric myoelectrical activity during different cognitive and physical tasks, including reading, doing bench work experiments, writing, and eating.

Fig. 7(b) presents the evolution of EGG signal amplitude (blue, left y-axis) and dominant frequency (red, right y-axis) over the 12-hour monitoring period, segmented into three representative activity states: *Reading, Doing Bench W*, and *Writing*. The amplitude reflects the strength of gastric slow wave activity, while the dominant frequency (in cycles per minute,) represents the intrinsic rhythm of gastric electrical pacing. During the experimental phase, both amplitude and frequency increased, suggesting elevated gastric activity likely driven by heightened physical or cognitive engagement. In contrast, during reading and writing, the amplitude remained lower and the frequency more stable. These observations indicate that EGG signals are sensitive to physiological states and mental workload, underscoring the potential of long-term EGG monitoring for a non-invasive assessment of gut–brain interactions.

### C. Discussion

The results demonstrate that FlexEGG, leveraging the AD8232 analog front-end (AFE) and the nRF52840 system-on-chip (SoC), is capable of acquiring high-quality EGG and ECG signals in a flexible, wearable form factor. The integration of these components enables a balance between signal fidelity, power efficiency, and wireless connectivity, which are the key requirements for continuous ambulatory monitoring. The use of a flexible substrate enhances user comfort and helps reduce motion artifacts compared to rigid devices. However, a quantitative comparison of motion artifact suppression with other flexible designs, such as those using textile electrodes, remains a valuable direction for future study. The low noise floor and high resolution of the AD8232 contribute to the clarity of the recorded ECG signals, suggesting that FlexEGG may be suitable for diagnostic applications, pending further clinical validation. The performance of nRF52840 in bio-potential testing has already been proved by previous research[14], and it also provides a robust Bluetooth Low Energy (BLE) link for wireless data transmission. Its low-power operation is essential for achieving practical battery life in continuous-wear scenarios, making it suitable for long-term physiological monitoring and provides insights which are difficult to obtain from short-duration tests. The continuous recordings allow the detection of transient arrhythmias, evaluation of treatment response and side effects, and generation of rich datasets for AI-based diagnostics, also reveals circadian patterns of gastric motility, enables detection of episodic dysrhythmias, and supports chronic disease tracking in conditions like gastroparesis or Parkinson’s-related dysmotility[11], [70]–[74]. Nonetheless, several limitations of FlexEGG remain. The current battery capacity may not support extended operation under advanced functionalities (e.g., on-board medical AI processing). In addition, larger-scale clinical trials are required to validate the device’s performance across diverse patient populations. Variability in skin types and the long-term adhesion stability of the flexible electrodes also warrant further investigation. Future work will focus on further miniaturizing the device to improve wearability and user comfort. The development of on-board signal processing algorithms for real-time event detection is also underway. We additionally plan to integrate complementary sensors, such as temperature and motion, to enable multi-modal physiological monitoring, and to explore alternative flexible materials and electrode configurations to further enhance comfort and long-term signal stability.

## IV. CONCLUSION

This paper presented FlexEGG, a flexible, patch-type wearable sensor designed for continuous, high-quality electrogastrography (EGG) monitoring, with simultaneous electrocardiography (ECG) acquisition. FlexEGG features a soft, stretchable, and skin-conformal electrode array composed of a hybrid conductive material, which ensures stable signal acquisition, low skin impedance, and enhanced user comfort during daily activities. By integrating the AD8232 analog front-end and the nRF52840 wireless SoC, the system provides an unobtrusive and reliable platform for long-term physiological monitoring. Preliminary results demonstrate high signal quality, robust wireless performance, and practical power consumption in real-world conditions. With further optimization and clinical validation, FlexEGG has a strong potential to advance non-invasive gastrointestinal health assessments and support the early detection and management of functional digestive disorders.

## Acknowledgment

J. L. thanks the support from the National Science Foundation under Award Nos. ECCS-2334134, ECCS-2216131, ECCS 2339495, EFMA-2318057, and CMMI 2323917. The authors thank Hang Yuan and Khoi Nguyen for assistance with the experiments.

