## Supplementary information for "Skin-conformal electronics for wearable electrogastrography monitoring"

Liuxi Xing, *member, IEEE*, Yulu Cai, Yunnuo Zhang, Vittorio Mottini, Linux Heller, Jinxing Li, *member, IEEE*

**Method**

**Morphological analysis (SEM):** The surface topography of the composite electrodes was examined using a JEOL 6610LV scanning electron microscope operated at an accelerating voltage of 15 kV.

**Mechanical testing:** Uniaxial tensile tests were performed on both freestanding films (≈100 µm thick) and films supported on elastic substrates using a CellScale UniVert equipped with a 20 N load cell. Rectangular specimens (30 × 10 × 0.1 $mm^3$) were gripped and elongated at a constant strain rate of 1%/s. Dimensions were determined with a caliper and validated under an optical microscope.

**Resistance under strain:** To assess electromechanical coupling, strips (10 × 25 × 0.1 $mm^3$) were mounted on the UniVert and electrically connected with copper leads bonded by silver epoxy. The samples were biased at 0.5 V, and resistance changes were monitored using a PalmSens4 potentiostat during stretching.

**Electrochemical impedance spectroscopy (EIS):** Impedance spectra were collected in phosphate-buffered saline (PBS, pH 7.4) over 1 Hz–1 MHz with a 10 mV sinusoidal perturbation, employing a PalmSens4 potentiostat.

**Conductivity measurements:** The bulk electrical conductivity of freestanding films was determined by a standard four-point probe method (Keithley 2400 source meter, room temperature). Thickness values were obtained with a micrometer and corroborated by optical microscopy.

**Thermal and humidity stability:** The temperature dependence of electrode impedance was evaluated in PBS solution between 23 and 60 °C using a three-electrode cell (sample as working electrode, Ag/AgCl reference, and glassy carbon counter). For humidity response, impedance was measured in a two-electrode configuration. All environmental stability tests were conducted using a Gamry Interface electrochemical workstation.

**Cytotoxicity test**

*Cell culture*: The immortalized human keratinocytes (HaCaT cells) were cultured in low calcium media made from no calcium DMEM (21068028, ThermoFisher Scientific), 10% FBS, and 1% penicillin-streptomycin at 37°C supplied with 5% $CO_2$ to maintain the dividing state.

*Sample preparation*: Followed a previously established protocol for a live-dead assay[1]. Briefly, FlexEGG electrode was placed in individual wells of a 96-well plate. 100 µl of a 1µg/ml fibronectin solution was added to each well and incubated at 37 °C for three hours. After the incubation period was completed, the excess fibronectin was removed from the wells, and each well was washed twice with 100 µl of sterile distilled water and once with 100 µl of cell culture media. An 80% confluent T75 flask of HaCaT cells was passaged, and a suspension containing 200 µl of low calcium media and 50000 cells was seeded in each well. The 96-well plate was incubated at 37 °C overnight.

*Stain solution*: The following day, a staining solution was prepared by mixing 2 µl of Hoechst, 5 µl of Calcein AM, and 10 µl of propidium iodide with 113 µl of low calcium media. 10 µl of the stain solution was added to each well and was incubated for 5 minutes at 37 °C.

*Imaging*: Samples were imaged using a Leica Thunder Imagers microscope with a 5x objective lens in brightfield and fluorescence channels for each stain.

*Cytotoxicity Assay Procedure*: To assess the biocompatibility of the electrode, a 24 hour live/dead cytotoxicity assay was performed. For individual well, fluorescence microscopy images were acquired for three channels: DAPI (labels all nuclei), Calcein-AM (stains live cells), and Ethidium homodimer-1 / Rhodamine (stains dead cells). Cell counts were extracted using CellProfiler (Broad Institute), which identified and quantified cells across all three fluorescence channels. Cell viability was calculated using the formula:

$$Viability\ (\%) = 100 - \frac{Dead\ cells}{All\ cells} \times 100 \qquad (1)$$

Control samples were intended to provide a baseline for spontaneous cell death in culture.

*Data Analysis*: Images were analyzed using a custom Cellprofiler workflow to count the number of cells in each channel and calculate cell viability. (https://github.com/YangLabUNL/PSEP-TEEI)

The safety of the FlexEGG electrodes through both in vitro cytotoxicity testing with human keratinocytes and in vivo skin irritation assessments in healthy volunteer were examined. As summarized in Fig. S1, keratinocyte cultures on the composite films exhibited high viability (96.52%) with negligible cell death, while human wear tests revealed no visible erythema, rash, or persistent skin marks after 24 h of continuous wear. These results confirm that the FlexEGG electrodes are non-toxic, non-irritating, and well suited for prolonged epidermal monitoring.

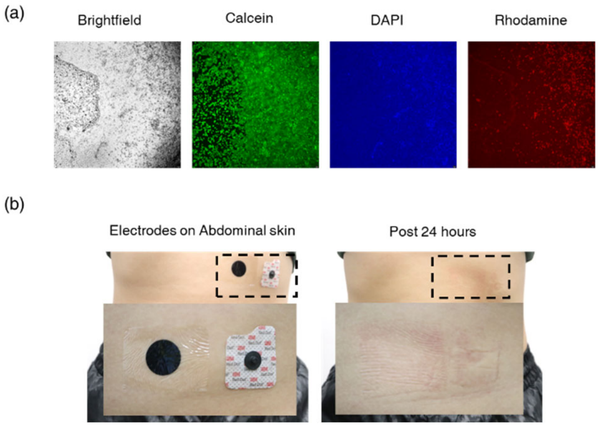


**Fig. S1** Biocompatibility of FlexEGG electrodes.
(a) Cytotoxicity evaluation with human keratinocytes cultured on representative composite films. Bright-field and fluorescence images (Calcein-AM, green; DAPI, blue; PI, red) confirm high cell viability (>95%) with negligible cell death, demonstrating minimal cytotoxicity of the electrode materials.
(b) Human Abdominal skin irritation tests comparing FlexEGG electrodes with commercial 3M Red Dot$^{TM}$ electrodes after 24 h of continuous wear. The FlexEGG electrodes showed no visible erythema or rash and skin marks disappeared within 1-2 h post-removal.

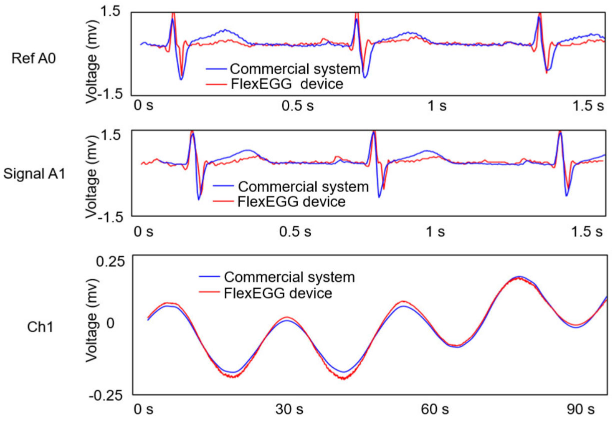


Fig. S2 The comparison experiment between proposed FlexEGG system and the commercial wearable holter.
Using the same setup as Figure 5 and 6, the raw EGG signals (channel A0, A1, and Ch1) from two systems are compared. It is seen that the proposed FlexEGG system provide similar result with ideal biopotential characteristic.

### Acknowledgment

J. L. thanks support from the National Science Foundation under Award Nos. ECCS-2334134, ECCS-2216131, ECCS 2339495, EFMA-2318057, and CMMI 2323917.

### References

[S1] S. R. Lorenzen, J. R. Brooks, T. C. Heiman, and R. Yang, “Porous substrate-based electroporation with transepithelial electrical impedance monitoring,” *J. Vis. Exp.*, no. 211, Sept. 2024.
